# Fishexplorer: A multimodal cellular atlas platform for neuronal circuit dissection in larval zebrafish

**DOI:** 10.1101/2025.07.14.664689

**Authors:** Sumit K. Vohra, Maren Eberle, Jonathan Boulanger-Weill, Mariela D. Petkova, Gregor F. P. Schuhknecht, Kristian J. Herrera, Florian Kämpf, Virginia M. S. Ruetten, Jeff W. Lichtman, Florian Engert, Owen Randlett, Armin Bahl, Yasuko Isoe, Hans-Christian Hege, Daniel Baum

**Affiliations:** Department of Visual and Data-Centric Computing, Zuse Institute Berlin, Takustraße 7, 14195 Berlin, Germany; Department of Computer Science and Engineering, New York University, NY, USA; Department of Molecular and Cellular Biology, Faculty of Arts and Sciences, Harvard University, Cambridge, Massachusetts, 02138, USA; Sorbonne Université, INSERM, CNRS, Institut de la Vision, F-75012 Paris, France; Centre for the Advanced Study of Collective Behaviour, University of Konstanz, Konstanz, Germany; Janelia Research Campus, HHMI, Ashburn, Virginia, USA; Laboratoire MeLiS, Université Claude Bernard Lyon 1 - CNRS UMR5284 - Inserm U1314, Institut NeuroMyoGène, Faculté de Médecine et de Pharmacie, 8 Avenue Rockefeller, 69008 Lyon, France

## Abstract

Understanding how neural circuits give rise to behavior requires comprehensive knowledge of neuronal morphology, connectivity, and function. Atlas platforms play a critical role in enabling the visualization, exploration, and dissemination of such information. Here, we present FishExplorer, an interactive and expandable community platform designed to integrate and analyze multimodal brain data from larval zebrafish. FishExplorer supports datasets acquired through light microscopy (LM), electron microscopy (EM), and X-ray imaging, all co-registered within a unified spatial coordinate system which enables seamless comparison of neuronal morphologies and synaptic connections. To further assist circuit analysis, FishExplorer includes a suite of tools for querying and visualizing connectivity at the whole-brain scale. By integrating data from recent large-scale EM reconstructions (presented in companion studies), FishExplorer enables researchers to validate circuit models, explore wiring principles, and generate new hypotheses. As a continuously evolving resource, FishExplorer is designed to facilitate collaborative discovery and serve the growing needs of the teleost neuroscience community.

## Introduction

A central goal of modern neuroscience is to understand how cellular and circuit-level mechanisms give rise to adaptive behaviors. Achieving this goal requires tools capable of bridging multiple spatial and temporal scales of neural data – linking synaptic connections to whole-brain networks, and transient neural activity to long-term developmental processes. Central to this challenge are neurons – diverse in type, intricately interconnected, and organized into functional circuits that span these scales.

The zebrafish (*Danio rerio*) has emerged as a powerful vertebrate model system for addressing this challenge, offering a rare combination of behavioral richness, genetic accessibility, optical transparency, and biological relevance even to the human brain. Their transparency enables real-time, *in vivo* brain-wide imaging at cellular resolution, while a sophisticated genetic toolkit – including CRISPR/Cas9-based genome editing, cell-type-specific activity reporters, and optogenetics tools – enable monitoring and perturbation of defined neuronal populations. Remarkably, despite their compact size, zebrafish larvae possess approximately 100,000 neurons (Bruzzone *et al.,* 2021), presenting an ideal balance between experimental tractability and circuit complexity.

A critical step in understanding neural circuits is validating connectivity. Traditionally, this has relied on physiological approaches such as electrophysiology or anterograde/retrograde labeling. Recent advances in serial electron microscopy now enable connectome-based validation through volume reconstruction at synaptic resolution. Notably, large-scale efforts like FlyWire (Dorkenwald, *et al.,* 2022, 2024) have demonstrated that whole-brain connectomics in *Drosophila* can reveal and predict the function of neural circuits underlying behavior. In vertebrates, partial reconstructions of the mammalian and human cortex (Shapson-Coe *et al.,* 2024, The MICrONS Consortium., 2025) have begun to uncover fundamental principles of neural computation and cell-type classification based on synaptic connectivity and morphology. In zebrafish, several groups (e.g., Svara *et al.,* 2022) are actively working toward a whole-brain connectome. In this context, Petkova *et al.*, 2025 captured ∼180,000 segmented soma and annotated ∼30 million synapses across the entire larval zebrafish brain (whole-brain EM dataset or WBEM), providing a comprehensive view of its neural architecture. Complementing this, Boulanger-Weill *et al., 2025* applied functional correlative light and EM (FCLEM) to link functional imaging data with ultrastructural reconstructions, enabling the identification of functionally and morphologically defined cell types and the exploration of circuit computation.

As multimodal datasets continue to grow in number and diversity – from calcium imaging to EM-based connectomics – there is a pressing need for platforms that meet three essential feature requirements: (1) spatial registration frameworks that align multimodal datasets within a standardized coordinate space; (2) integrated neuron databases that consolidate key features of individual cells, including morphology, molecular identity, functional tuning, and genetic labels – enabling structured queries and circuit-level analyses; and (3) native visualization and data analysis tools that provide 3D rendering, exploration across imaging modalities, as well as attribute- and similarity-based neuron searches to support data-driven hypothesis generation about neural circuits. While existing atlases implement subsets of these features, no current resource combines all three within a unified, interactive framework.

Over the past decade, the zebrafish neuroscience community has produced a suite of complementary brain atlases that have enabled multiscale investigations – spanning anatomical, genetic, functional, and behavioral datasets – though often in a siloed manner (Légaré, et al., 2023, Marquart, et al., 2017). In larval zebrafish, the Z-Brain Atlas provides a foundational 3D anatomical reference for 6-day-old larvae, segmenting the brain into 294 regions using antibody-based registration techniques (Randlett et al., 2015). This atlas has become central to transcriptional analysis and functional imaging studies, enabling the spatial localization of signals within a common coordinate framework (Lange et al., 2024). Also, the Adult Zebrafish Brain Atlas (AZBA) has addressed long-standing imaging challenges through advanced tissue-clearing protocols and light-sheet microscopy (Kenny, et al., 2021). With 217 segmented regions and integration of ten immunostains and nuclear markers, AZBA provides the first comprehensive reference of the adult zebrafish brain. This atlas has enabled automated segmentation of brain-wide gene expression patterns, revealing major conservation in neuronal subtypes between larval and adult stages. Building on this foundation, mapZebrain introduced cellular-resolution morphological reconstructions from LM data by mapping over 4,300 individual neurons into a standardized larval brain template (Kunst et al., 2019), although axons and dendrites are indistinguishable. But, this resource integrates transcriptomic profiles (Shainer et al., 2025), offering a rich multimodal view of neuronal identity and projection.

Despite recent progress, critical limitations remain. First, platform fragmentation hinders interoperability – datasets remain scattered across incompatible systems, complicating integration. Second, most atlases are static and lack community-driven extensibility, with limited support for user uploads, APIs, or third-party tool integration, leaving no open framework that enables researchers to analyze their data alongside others’. Third, multiscale integration remains incomplete: synapse-level connectivity is rarely linked to cellular or circuit-level features. Fourth, structural data remains siloed from dynamic functional modalities like calcium imaging or behavioral annotations. Fifth, search functionality is weak – no platform supports complex, attribute-based queries across modalities.

Here, we present the **FishExplorer** platform (https://fishexplorer.zib.de/sandbox/), which directly addresses these gaps and the aforementioned requirements through its integrated architecture. FishExplorer not only streamlines analysis workflows but also enhances reproducibility with standardized data formats and fosters collaboration via a shared, interactive annotation environment. FishExplorer is a web-based, multimodal, and multiscale neuroanatomical atlas designed to meet the growing demands of zebrafish neuroscience.

FishExplorer is built around the FAIR principles (Wilkinson et al., 2016): it ensures data are findable (via rich metadata and UUID-based identifiers), accessible (with clear access protocols and tiered user permissions), interoperable (using standardized formats and compatible with external tools), and reusable (through open licensing and detailed documentation). FishExplorer supports standard data formats and naming conventions to ensure that both internal and external datasets remain accessible and interoperable – an essential feature for collaborative and reproducible research. By employing manual and automated registration tools (Avant et al., 2007, Bogovic et al., 2016), FishExplorer fosters the integration of diverse imaging modalities – including LM, EM, X Ray micro-computed tomography (CT) – into a shared coordinate framework anchored by a standardized larval reference brain (Randlett et al., 2015). This interoperability ensures compatibility with a range of widely used zebrafish neuroscience tools and enables continuity, reuse, and expansion of existing work. Integration with the EM reconstruction tools CAVE (Connectome Annotation Versioning Engine) (Dorkenwald et al., 2025) and Neuroglancer allows users to view and query synapse-resolved connectivity while keeping them contextually embedded in whole-brain LM maps. This bridge between LM and EM is particularly powerful for integrating cell-type properties, discovering connectivity motifs, and correlating activity patterns with fine structural detail. FishExplorer hosts a rich, queryable, diverse neuron database that consolidates cell-level features such as morphology, neurochemical identity, connectivity, and function. It supports similarity-based neuron comparisons using custom NBLAST matrices tailored for zebrafish and offers clustering tools for exploring potential cell types or conserved morphologies. A suite of interactive visualization and analysis tools allows users to render brain structures and neurons in 2D and 3D, overlay synapse positions, and compare circuit structures across datasets. FishExplorer also implements a tool (Vohra et., 2024), that enables intuitive hypothesis generation through visual search and comparison, with filtering across anatomical regions, functional attributes, and metadata tags.

In summary, FishExplorer unifies anatomical maps, neuronal databases, and analysis tools within a single interface. FishExplorer is designed for growth: new data can be uploaded post-publication, aligned with existing content, and made available to the broader community – encouraging transparent, collaborative discovery.

## Results

### FishExplorer platform

Having outlined the motivation and scope of FishExplorer, we now present the platform’s architecture and core features, demonstrate its interoperability with different imaging techniques and datasets, and show how researchers can use the interactive tools to explore, compare and share data in a standardized way and achieve reproducible results. As illustrated in **Fig. 1a,** FishExplorer unifies anatomical reference brains, neuronal databases, and analysis tools within a single web-based interface. Furthermore, FishExplorer enables researchers to integrate, explore, and share their own data as well as perform brain-wide structural analysis of neuronal morphologies. The platform’s interoperability is demonstrated in the “Brain regions and bridges” and “Neuron database” sections, which describe the integration of multiple light and electron microscopy datasets for the larval zebrafish. To make the data findable and reusable, the platform is equipped with a set of analysis tools that allow users to interactively create search queries, collect relevant neurons from different databases, and compare them. This functionality is demonstrated in sections “Custom NBLAST Matrix and Neuron Classification” and “Search, exploration and comparison across databases”. Regarding accessibility, all data stored in the platform can be downloaded, and users can share their own data by uploading it, thereby making it accessible to other researchers. This is demonstrated in section “Community engagement and data enrichment for FishExplorer”.

**Fig. 1.**
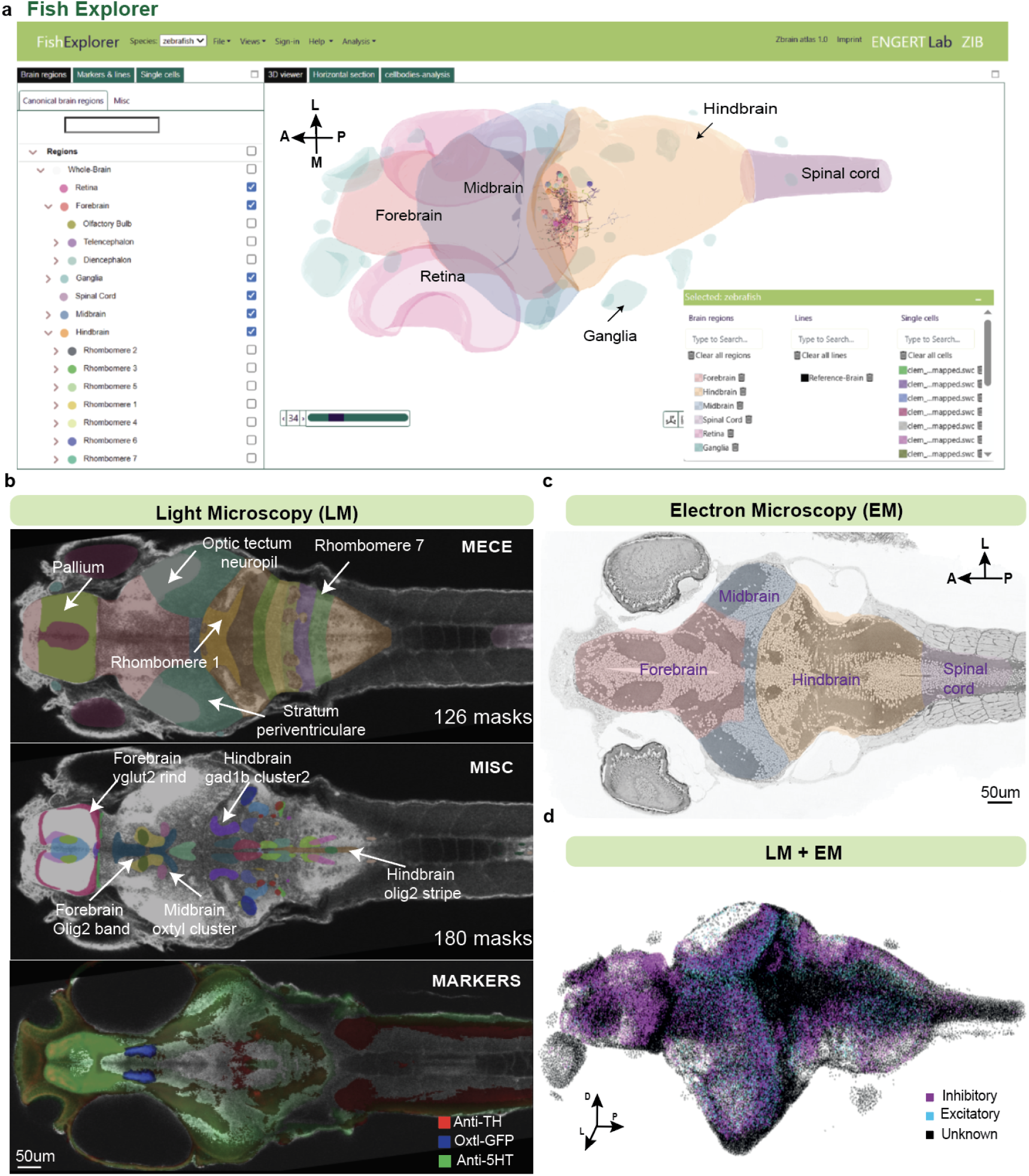
FishExplorer and multimodal information for the larval zebrafish. **a**, Overview of FishExplorer. FishExplorer is an interactive web-based atlas platform that enables users to select anatomical region masks from brain to body and visualise multimodal datasets in both 2D and 3D. On the left side of the window, regions of interest can be selected from the ‘Brain regions’ tab; single cells of interest can be selected from the ‘Single cells’ tab. On the right bottom of the window, selected items are listed in a separate window, where users can change the items’ color. FCLEM cells are shown with MECE masks as an example. **b**, Region masks, such as MECE (top) and MISC (middle) masks, were segmented in LM data, shown here in a 2D representation; Transgenic lines, molecular markers, and EM datasets can be selected via the “Markers & Lines” tab in (**a**). An example overlay shows three different markers on the reference brain (bottom). **c**, MECE masks overlay on a slice of the WBEM stack. **d**, 3D segmented cell bodies from WBEM with cell type labels, defined by neurotransmitter expression (purple: gad1b expression, inhibitory neuron; blue: vglut2a expression, excitatory neuron), are shown.

### Brain regions and bridges

To elucidate the neural connection and understand circuit function, anatomical information is critical. On the platform, users can interactively select anatomical regions masks from the defined ‘brain regions’ and visualize them (**Fig. 1a, b**). The brain regions are divided into two categories: MECE (Mutually Exclusive and Comprehensively Exhaustive) and MISC (Miscellaneous) regions. MECE masks cover the whole brain and are organized hierarchically. They provide a comprehensive, non-overlapping representation of brain regions that allows systematic and quantitative analysis. The MISC regions complement the MECE regions and are used for targeted sub-populational analysis, so they are less strictly defined and can overlap, such as neural clusters that express specific neurotransmitters. They were defined based on several marker-gene expressions, such as neurotransmitters and neuropeptides. In total, 126 MECE and 180 MISC masks were defined for the larval zebrafish. We also incorporated the autonomic nervous system into our catalog of brain region masks, allowing integration of information about organs and nerves outside the brain (**Extended Data Fig. 1a, b**). Additionally, users can select and visualize slices of fluorescent, X-ray and calcium imaging stacks and overlay them (**Fig. 1c**).

To map and transfer information between different modalities, we generated several transformational bridges, such as LM-EM bridge (Petkova *et al., 2025*; Boulanger-Weill *et al.,* 2025) and LM-X-Ray bridge (**Methods**). Using the LM-EM bridges allowed us to map MECE and MISC masks onto the low-resolution WBEM and FCLEM datasets, respectively (**Fig. 1a)**.

### Neuron database

Neuron datasets are the key resource that allow neurobiologists to localize neural circuits involved in the emergence of specific behaviors. These datasets are typically generated using diverse light and serial-section electron microscopy imaging techniques, each with its own advantages and limitations. Several neuron datasets for zebrafish – both established and more recent ones – have been created by different research groups (e.g. mapZebrain (Kunst *et al.,* 2019), EM datasets (Hildebrand *et al.,* 2017, Petkova *et al., 2025, &* Boulanger-Weill *et al., 2025*), and photoactivatable green fluorescent protein (PA-GFP) (Boulanger-Weill *et al.*). However, no universal standard for naming these datasets exist. Here, we have implemented the following naming convention in our platform: lab name, dataset name, and year of creation. To ensure that neurons remain discoverable and reusable in the future, we have implemented a comprehensive metadata annotation mechanism. Neurons are annotated with the list of meta-tags e.g., tracer’s name (**Extended Data Table 1**). This metadata is incorporated at the top of the SWC file. To enable comparison and cross-linking of these datasets, we registered them to our common reference brain (Randlett *et al.,* 2015).

Currently, FishExplorer incorporates four zebrafish neuronal datasets – WBEM (Petkova *et al.*), FCLEM and PA-GFP (Boulanger-Weill *et al.,*), and mapZebrain (Kunst *et al.,* 2019). In total, we integrated 4600 neurons into the platform (WBEM: 1200, FCLEM: 241, PA-GFP: 47, mapZebrain: 3120) (**Fig. 2b**). The table in **Fig. 2a** summarizes all the important aspects of these datasets. For the WBEM dataset, serial-section electron and light microscopy techniques were combined (Petkova *et al.*) to map brain structure in the whole brain of the same animal with cell-type information (**Fig. 2c**), whereas FCLEM is a hypothesis-driven dataset designed to map the structure-function relationship of each neuron in the anterior hindbrain brain (**Fig. 2d**). In the WBEM and FCLEM datasets, axons and dendrites are distinguishable. The PA-GFP (**Fig. 2e)** consists of light microscopy-based neuron morphologies (Boulanger-Weill *et al.*) reconstructed in different individuals at 7dpf in the anterior hindbrain. mapZebrain is a sparse connectome, comprising light microscopy-based neuron morphologies reconstructed by Baier and colleagues (Kunst *et al.,* 2019) in different individuals of approximately the same age (5-7 dpf; **Fig. 2f**). Reconstructed neurons were sparsely labeled and consist of approximately 4,300 neurons. Meta information is available for all datasets except mapZebrain and is included in the header of the SWC files.

**Fig. 2.**
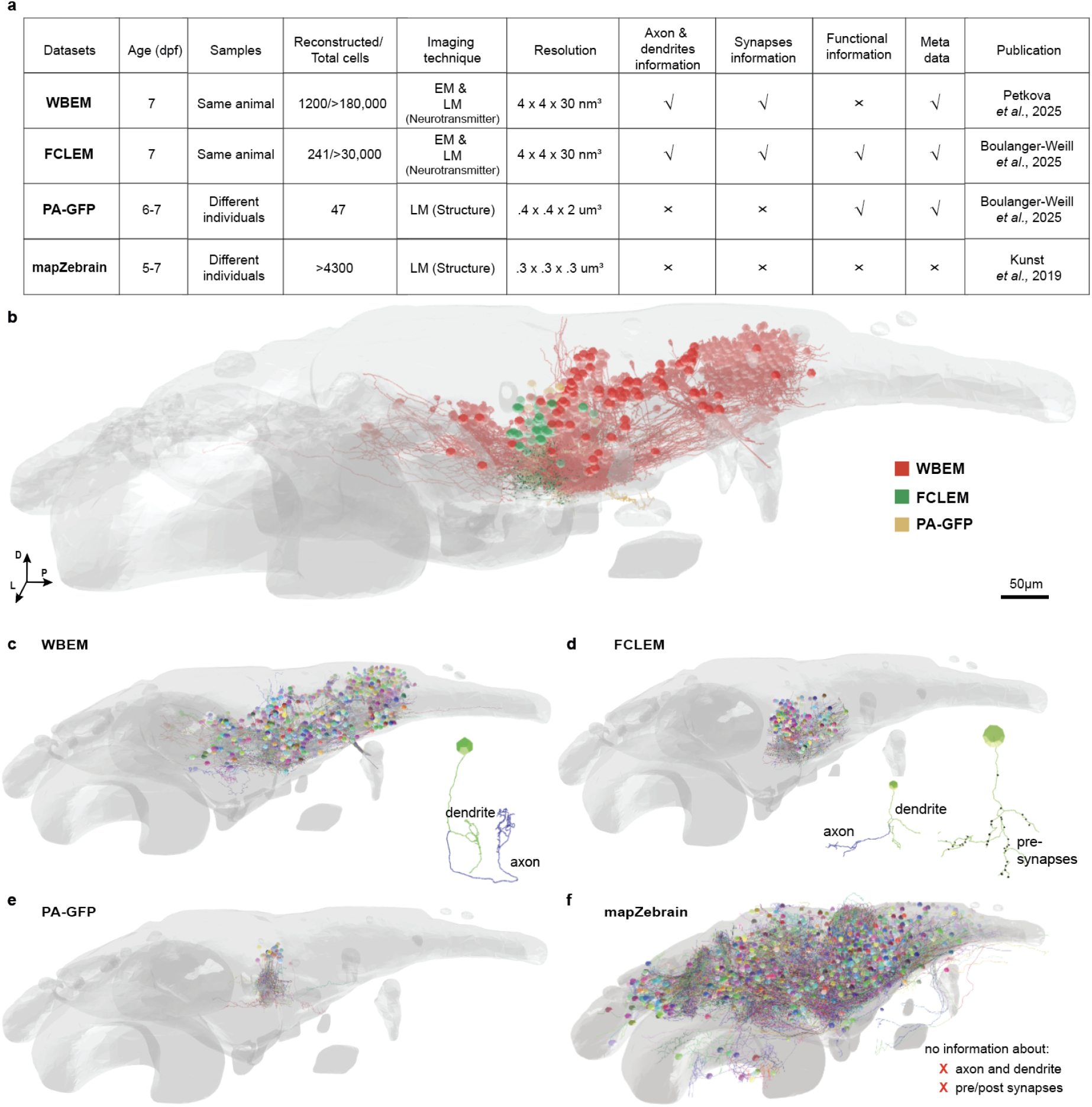
Neuron database. **a**, Overview of the cellular datasets integrated in FishExplorer. In the ‘Imaging technique’ column, ‘LM (Neurotransmitter)’ indicates gene expressions were observed with LM across the whole brain, and ‘LM (Structure)’ indicates that cellular morphologies were identified with LM by GFP expression in subsets of neurons. **b**, Overview of integrated datasets: newly reconstructed neurons in the companion papers integrated into the FishExplorer are shown, depicting the diversity and coverage of neuronal morphologies. Colors of neurons are assigned based on the dataset as labels show. **c, d**: EM neuron reconstructions. **c**, Reconstructed neurons from a WBEM dataset, including axons and dendrites information. **d**, Reconstructed from a FCLEM dataset, with axons and dendrites information, and pre-/post-synapses (black circles indicate synaptic location). **e, f,** LM neuron reconstructions:Reconstructed from a PA-GFP dataset, labeling by photo-activation and mapped via our registration pipeline (**Fig. 5a**). **f**, mapZebrain neurons mapped via our registration pipeline (**Fig. 5b**). This dataset doesn’t include axons and dendrites information, and pre-/post-synaptic information.

### Custom NBLAST Matrix and Neuron Classification

Organization of large-scale neuronal databases remains a significant challenge, and researchers commonly classify neurons based on position of the soma and morphological similarity. One widely used tool for this purpose is NBLAST (Costa *et al.,* 2016), which enables pairwise comparison of neuronal shapes based on their geometric features. NBLAST is commonly used for the Drosophila brain, e.g., in Schlegel *et al.,* 2024 where a new definition of ‘cell types’ was proposed based on comparative analysis of morphologies across three brain hemispheres). A core entity of NBLAST is the precomputed scoring matrix, which encodes the log probability ratio between the matching and nonmatching segments. The score for a neuron pair depends on the scores between their segments. Based on the joint distributions of distance and absolute dot product between segments, the probability of matching and nonmatching segments is derived. This is used to construct a scoring matrix which describes a segment score (in Euclidean space) based on the dot product (rows) and distance in micrometers (columns). The size of the matrix is arbitrary; we chose a 10×21 array, as for the fly matrix. However, similarity scores are sensitive to factors such as the model organism (i.e., the volume from which neurons are traced) and the imaging modality (e.g., LM vs. EM, or neurons reconstructed using resampling techniques). The default matrix was originally trained on Drosophila LM data, which may limit its performance when applied to other datasets. To address this, we extended the existing approach to compute a custom matrix using a combined dataset from the mapZebrain (Kunst *et al.,* 2019) and our own FCLEM zebrafish neurons, and compared it with the default matrix (**Fig. 3a, left**). The methodology for matrix construction based on multiple datasets is detailed in the **Methods**.

**Fig. 3.**
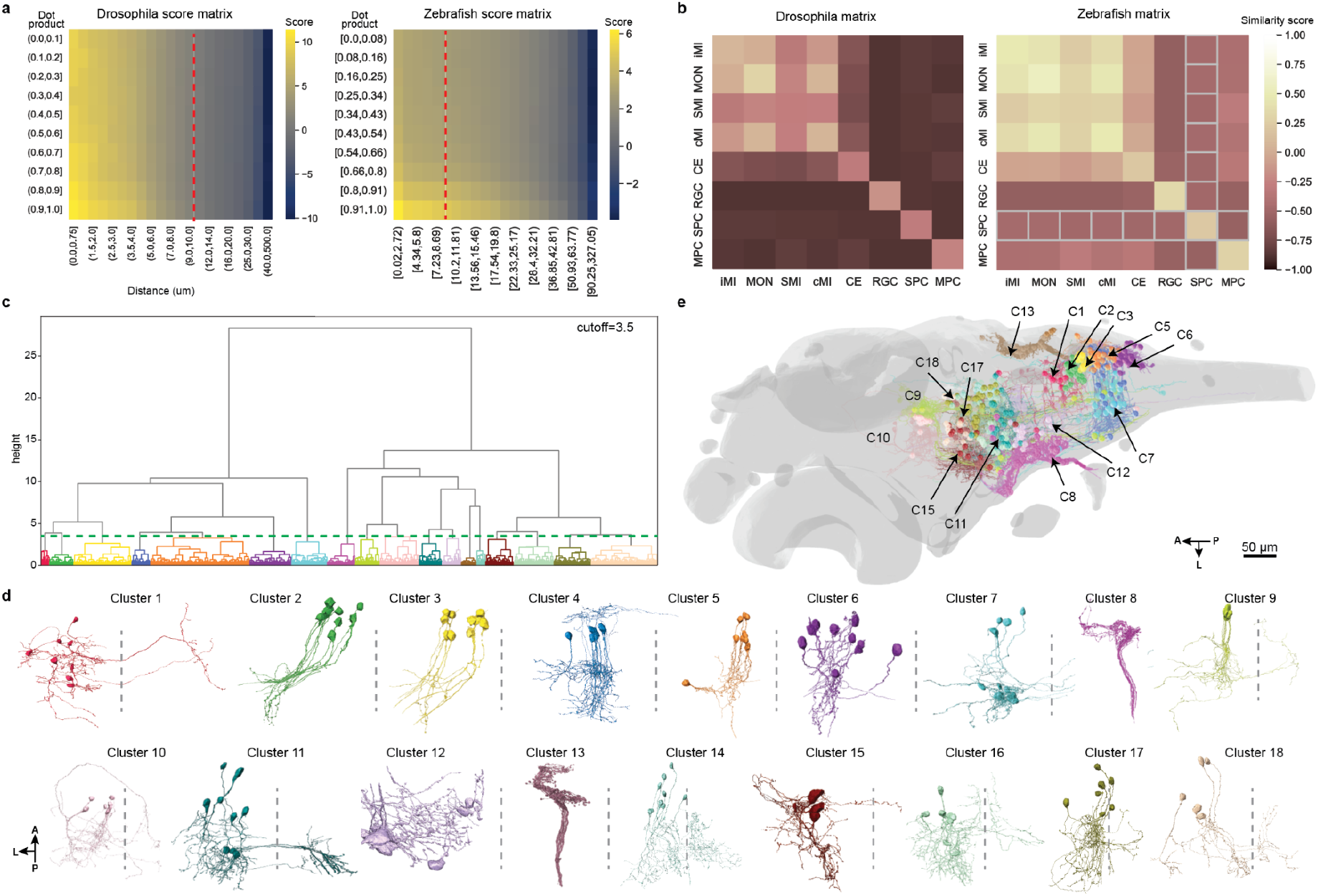
Comparison of fly vs zebrafish NBLAST score matrices and clustering of neuron datasets. **a,** Matrix comparison and validation: The default NBLAST score matrix (left), originally derived from Drosophila LM data, is compared with a custom score matrix (right) trained on zebrafish datasets. Red dotted lines indicate the column where the distance is 10 um. **b,** Matrix comparison and validation: Both matrices were applied to a test dataset with known zebrafish cell types from FCLEM and mapZebrain (i.e., ipsilateral motion integrator (iMi), contralateral projecting motion integrator (CMI), motion onset (MON), slow motion integrator (SMI), cerebellum eurydenroid (CE), retinal ganglion cells (RGC), subpallium principal cells (SPC) and medulla principle cell (MPC)). The custom matrix (right) showed improved performance, reflected in higher average pairwise similarity scores within cell types. **c**, Hierarchical clustering of all WBEM and FCLEM neurons using the custom NBLAST matrix: The dendrogram was cut at the height indicated by the dashed green line (cutoff=3.5), resulting in 18 distinct clusters as colored in different colors. **d**, Cluster visualization: 3D representations of representative neurons (n=10 for each cluster) from the 18 identified clusters, highlighting their structural diversity and spatial distribution, shown in dorsal view. The dotted lines indicate the midline of the brain. **e**, s all the 18 clusters are mapped on the reference brain. Black arrows indicate the location of the somas of each cluster.

We present the NBLAST matrix with raw scores created from zebrafish neurons (**Fig. 3a, right**) and compare it to the scoring matrix created from Drosophila neurons (**Fig. 3a, left**). Notably, the zebrafish scoring matrix distinguishes longer distances than the fruit fly scoring matrix. This likely addresses difficulties in grouping distantly related neurons in the zebrafish with the default matrix described by Costa et al. (2016). For example, for the fruit fly scoring matrix, over 60% of columns describe distances up to 10 µm (indicated by red lines in **Fig. 3a**), while for the zebrafish, less than 35% of columns are in this range. Qualitatively, we observe a stronger dependence of the raw score on the dot product (rows) than on distance (columns) for the zebrafish scoring matrix. To evaluate how the new matrix affects similarity scores of zebrafish neurons, we calculated the NBLAST scores with both matrices for held-out neurons from the matching and non-matching groups (test set). A successful metric should provide high scores within matching neurons, and low scores across matching and non-matching, and within non-matching neurons. In **Fig. 3b**, scores across the neuron types from which matching neurons were chosen are provided. Neurons were selected from cell types determined by Kunst *et al.,* 2019 and Boulanger-Weill *et al.*, 2025 (**Methods**). Within all types, we observe an increase in the average NBLAST similarity score for the zebrafish over the fruit fly scoring matrix. Furthermore, we see a better distinction between types for the zebrafish scoring matrix. As described above, the fruit fly matrix assigns low scores for shorter distances than the zebrafish matrix, which can lead to low similarity assignments when neurons are larger, as is the case in the zebrafish brain.

Next, we used the zebrafish scoring matrix to cluster zebrafish neurons from the FCLEM neurons that are functionally labeled as hindbrain integrator circuit components (**Extended Data Fig. 3a, b**). The dendrogram shows the hierarchical relationship of similarity across the neurons. When cut at a height of 1.3, we confirmed four clusters corresponding to the neurons identified in the hindbrain integrator circuits. This automated classification achieved an F1 score of 69.2%, demonstrating reasonable accuracy. To improve classification performance, Boulanger-Weill *et al.*, 2025 first extracted a wide range of structural features of the neurons and then applied traditional machine-learning classifiers.

To further validate the approach, we applied the same custom matrix to cluster zebrafish neurons from the WBEM and FCLEM dataset (**Methods**). When cutting the dendrogram at a (dimensionless) height of 3.5, we identified 18 clusters (**Fig. 3c-d**). Several of these correspond to known cell types: cluster 1-5 matches to the motor vagus neurons; cluster 8 and 13 match PLL (posterior lateral line) cells; cluster 9 and 10 matches TEN (the tegmental excitatory neurons); cluster 12 corresponds to reticulospinal neuronal cell types while others appear to be more heterogeneous.

### Search, Exploration and Comparison Across Databases

Neuroscience research often involves hypothesis-driven exploration – for instance, investigating neuromodulatory pathways or circuit-level mechanisms underlying specific brain functions and sensorimotor transforms. To enable efficient search, exploration and comparison of neurons across multiple datasets, we integrated a visual analytics tool, the CircuitExplorer, into the FishExplorer platform. CircuitExplorer supports such investigations by providing an interactive, visually guiding interface that integrates data from various modalities, including microscopy and behavioral experiments.

The previous version of CircuitExplorer (Vohra *et al.,* 2024) offered only basic spatial filtering capabilities and was limited to a single neuron database. We have significantly extended its functionality by introducing meta filters, NBLAST-based similarity search, and support for cross-database comparisons. Computing similarity scores and filter queries dynamically across all available datasets is computationally intensive and memory-demanding. To address this, we precomputed similarity models and stored them in the backend database. These models are periodically updated as new neuron morphologies, anatomical regions, or additional metadata become available. Additionally, users can link their custom datasets with existing model sets on demand. The web interface of CircuitExplorer (**Fig. 4a**) includes several key components, designed to facilitate flexible query construction and comparative analysis: (1) The ‘Query Builder’ (**Fig. 4a, left**) features two juxtaposed tree structures, enabling users to build complex, logic-based queries using drag and drop functionality. Queries can target major neuron features (**Fig. 4b**), such as position, morphology, function, molecular markers, brain regions (MECE and MISC masks) and metadata (e.g., tracer name, publication source). The left subtree lists all available attributes in the form of set, spatial and meta operators necessary to build a query, while the right tree helps assemble query logic. Once a query is executed, (2) the ‘Pathway Browser’ (**Fig. 4a, center**) displays all candidate circuitry related to the selected features, i.e. possible pathways matching the current query. Finally, (3) the ‘Neuron Browser’ presents subsets of matching neurons and highlights structurally similar neurons using NBLAST-based comparisons. Additional panels for visualizing neuronal morphologies, synapse positions, and anatomical overlays are available, with usage guidelines provided in the platform’s online tutorials.

**Fig. 4.**
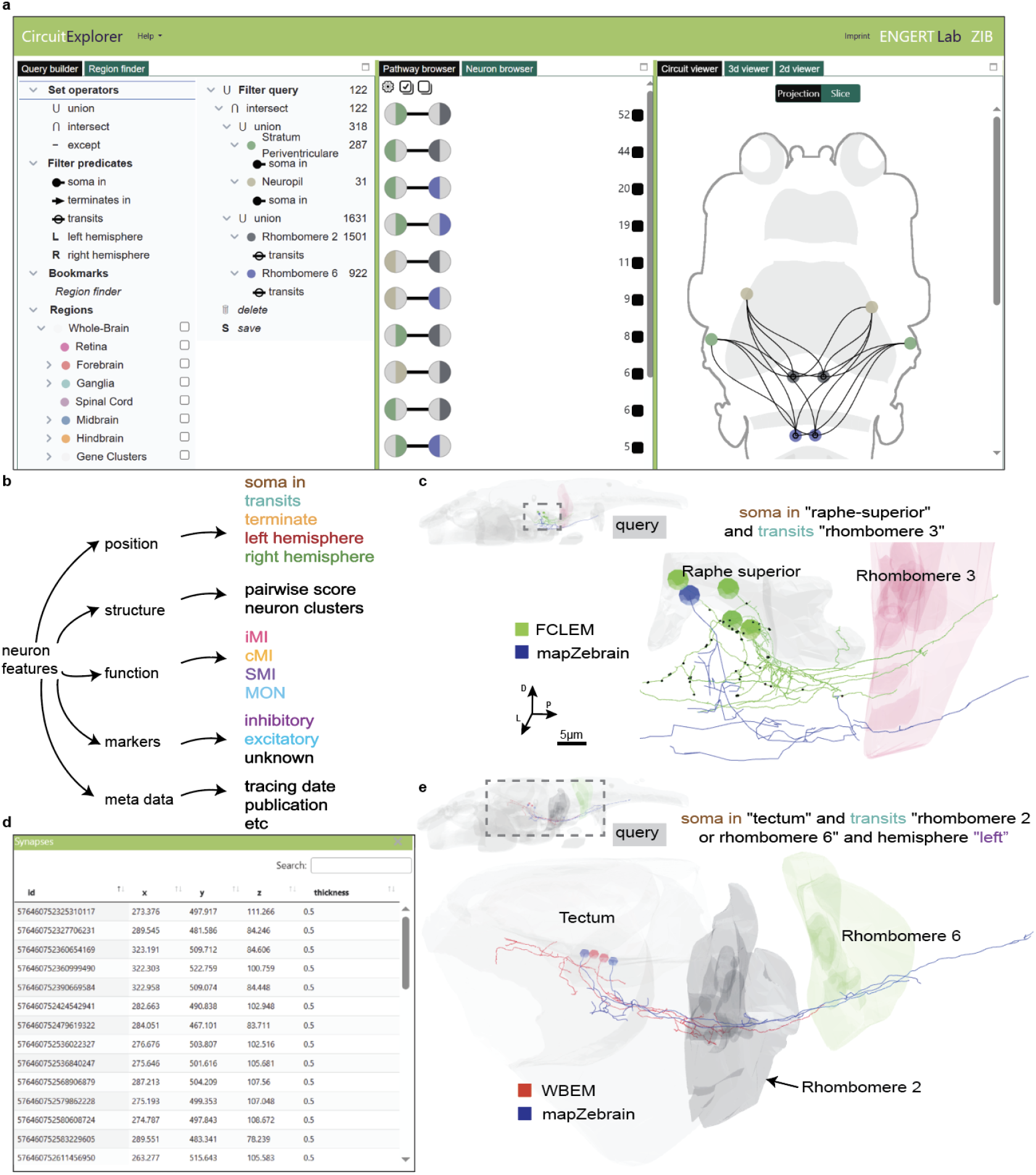
CircuitExplorer: Interactive search and cross-dataset linking of neurons. **a**, CircuitExplorer interface: Overview of the interactive web-based CircuitExplorer analysis tool. Users can visually explore and search for neurons of interest across multiple integrated datasets. An example query mentioned in **e** is shown. **b**, Search modalities: Neurons can be queried and linked across databases based on five major attributes: anatomical position, morphological feature, functional properties, molecular markers and associated metadata. **c**, Cross-dataset structural similarity search: Example query demonstrating how users can filter neurons having “soma in Raphe superior and transits through Rhombomere 3” in the mapZebrain dataset, and identify structurally similar neurons in the FCLEM dataset using NBLAST. **d**, Example demonstrating how users can visualize pre-post synapses (x, y, z represents the location of the particular synapse along with its thickness) information related to selected neurons from CAVE. **e**, Advanced filtering across datasets: Illustration of complex multi-attribute filtering applied to identify specific neuron subsets within the mapZebrain and WBEM datasets.

As an example to search neurons whose soma locates in a specific brain region and to visualize their morphologies, we show a query for “neurons having soma in the Raphe superior” which leads to examine specific serotonergic populations (Kawashima *et al,* 2016) and their circuit-level roles across datasets (**Fig. 4c**). First, we queried the mapZebrain dataset and found eight neurons whose soma are in the Raphe superior. Next, we selected a particular neuron (colored blue in **Fig. 4c**) and searched for similar neurons in the FCLEM dataset using the NBLAST-based forward score. We then visually compared neurons in an anatomical 3D viewer and identified four potentially similar neurons (colored green in **Fig. 4c**). To assess whether these two neurons belong to the serotonergic population, we loaded the serotonin (5-HT) stained line in the anatomical 3D viewer and checked for overlap. Since both cell somas are close to the boundary of the serotonergic cluster, we could estimate their serotonergic identity.

As a second example, we illustrate the power of CircuitExplorer to perform complex multi-attribute filtering, combining spatial, metadata, and marker features across datasets. Specifically, we searched for “Tectal neurons having soma in the left hemisphere and projected to either Rhombomere2 or Rhombomere6”. Such neurons have already been described (Sato et al., 2007) but their postsynaptic targets remain underscribed.

First, we queried the mapZebrain dataset and upon executing the query, we found 55 such neurons. Next, we selected neurons looking for neurons that stay in the same hemisphere and selected the top 2 (colored blue in **Fig. 4d**). Next, we searched for similar neurons in the WBEM dataset and also incorporated a meta filter in the existing query. On query execution, we found 2 such neurons (colored red in **Fig. 4d**). Furthermore, users can open these 2 neurons in the Neuroglancer or CAVE browser by clicking their links available in the metadata tab and trace their potential postsynaptic partners.

These two examples highlight how simple queries enable detailed circuit-level exploration and identification of potential postsynaptic partners contributing to sensory-motor processing.

### Community engagement and data enrichment using FishExplorer

The FishExplorer is designed to encourage community engagement and continuous enrichment with published data across teleost fish research. The process of incorporating user-defined brain regions, molecular markers and transgenic line datasets was introduced in a previous paper on the cross-species functionality of the atlas platform (Vohra *et al.,* 2024). Here, we describe two independent processes by which the users’ data can be visualized and integrated into the platform (**Fig. 5**).

**Fig. 5.**
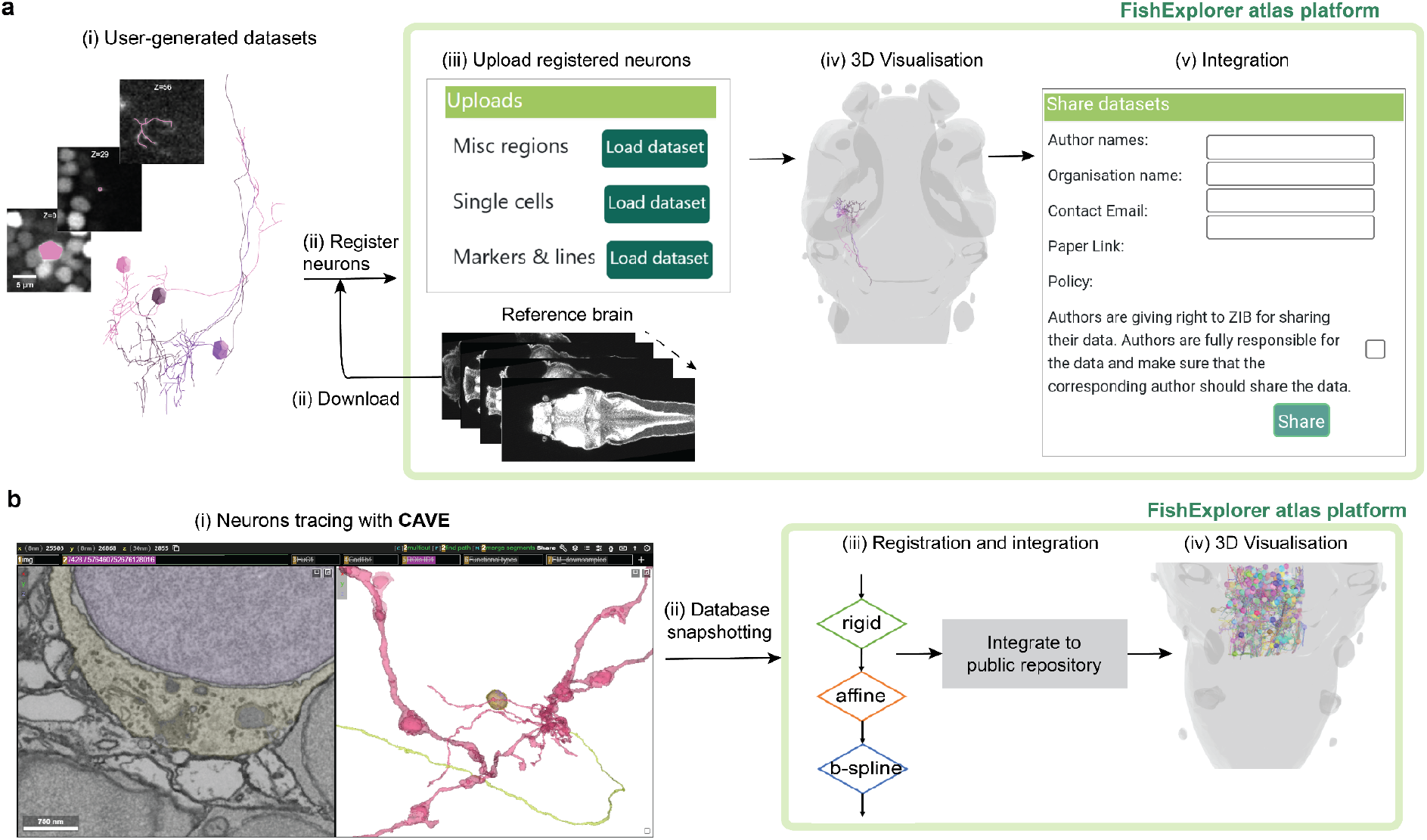
Neuron integration in FishExplorer. **a**, Step-by-step workflow for integrating user-generated datasets into the FishExplorer platform: (**i**) Here a few cells from the PA-GFP dataset are shown as an example. The photoactivated neurons were imaged in a confocal microscope and neuronal skeletons were reconstructed later. After preparing the custom LM neuronal dataset, (**ii**) users can download the reference stack and Python registration script for registering their dataset to the zebrafish reference brain. Registration is performed locally by the user. (**iii**) Once registered, users log in to the platform and upload their registered dataset. The uploaded neurons are then automatically processed and made accessible exclusively to the corresponding user. By default, the dataset remains private and will be visible only to the corresponding user. (**iv**) Users can then visualise/analyse their neurons alongside existing single-cell and brain region datasets within FishExplorer. (**v**) Upon publication of a paper using this dataset, and at the user’s request, the dataset can be added to the publicly accessible single-cell database within the platform. **b**, Automated workflow for incorporating WBEM and FCLEM neurons into FishExplorer: (**i**) Users can trace or update neurons in the whole brain electron microscopy and FCLEM datasets using CAVE client. The traced/updated neurons are stored in the database. (**ii**) Database snapshotting: At regular intervals, snapshots of the live database are created to preserve and manage versioned data. (**iii**) Registration and integration: The platform then automatically retrieves updated and newly traced neurons from each frozen snapshot and registers them to the zebrafish reference brain using ANTs-based rigid, affine and B-spline transformations. registered neurons are automatically added to the public directory of the “Single Cells” database. (**iv**) 3D visualization: Example 3D rendering of WBEM and FCLEM neurons within the zebrafish brain atlas.

First, we show the recipe to upload custom user single-cell datasets (**Fig. 5a**). (A detailed step-by-step tutorial to volume and point based registration is available at the **FishExplorer**website):

1. **Registration to the reference brain:** The user downloads the reference image stack and registration code from FishExplorer and registers the custom image stack to the reference stack outside the platform. Forward and inverse transformations are generated upon completion of the image registration. Next, the custom SWC neuron files (skeletons, consisting of points) are registered using the generated transformations.
2. **Upload and analysis:** The user can upload the registered neuron files to the platform and initially load them into the user’s private repository in FishExplorer. This allows users to visualize, link and compare their custom neuron morphologies with those in existing datasets.
3. **Community access:** Upon publication and user request, the datasets are transferred and integrated into FishExplorer’s public repository, making them accessible to the community.

Second, we also developed an automated process of integrating neurons that are segmented by the community for WBEM and FCLEM datasets into the platform (**Fig. 5b**). Here, users segment neurons of interest using CAVE (Dorkenwald *et al.,* 2025). CAVE uses a cost-efficient proofreading and segmentation approach called ChunkedGraph, which enhances collaborative reconstruction of neurons (Dorkenwald *et al.,* 2022). Since generating skeletons (sparse representation of neurons) and meshes from reconstructed neurons using ChunkedGraph is time-consuming, CAVE creates snapshots of these datasets at regular intervals for faster retrieval and analysis. The FishExplorer platform downloads these new or updated reconstructions from the latest snapshot, and registers and incorporates them into the platform (**Fig. 5b**).

## Discussion

We presented a comprehensive platform that allows users to visualize, search and analyse cell body locations, morphologies, connections and cell types across multiple datasets of zebrafish larvae. Traditionally, there was no central, open, and extensible atlas framework that allowed researchers to analyze their own data in the context of others’ – a process that remains time-consuming and dependent on individual collaborations. FishExplorer allows users to bridge across data types and scales, from whole-brain imaging to synapse-level anatomy, and to explore structure-function relationships across modalities.

The NBLAST tool allowed us to identify similar neurons and categorize them based on structural morphology. The incorporation of NBLAST-based comparisons inside the CircuitExplorer supports the analysis across datasets. Using a custom score matrix, we achieved a better matching score between query-target neurons as compared to the fly matrix, for which the matching score was always low. Precomputing the all-by-all neuron score matrix and storing the top 500 hits for each neuron enabled interactive real-time analysis on the web-based platform.

In the CircuitExplorer analysis tool, models are precomputed, which enables rapid searches and highly interactive complex multi-attribute filtering of neurons across the existing multiple datasets. The pathways generated by CircuitExplorer are purely based on spatial information, therefore the connections from ‘region A’→’region B’, ‘region B’→ ‘region C’, does not guarantee that there is an actual direct connection across ‘region A’→ ‘region B’→ ‘region C’, so researchers should validate the actual biological connection by the EM reconstruction and other labeling techniques. CircuitExplorer currently uses synapse information to a limited extent; however, we do not incorporate synapse information during query building, since the inclusion of ∼30 million synapses and linking it with the rest of data is a challenging task in terms of analysis and scalability.

FishExplorer is designed to be extended and expanded with multiscale datasets. In the near future, we will extend our platform with more anatomical regions from other body organs, such as autonomic nervous system, and other well known single-cell datasets from the teleost research community (Hildebrand *et al.,* 2017, Svara *et al.,* 2022). We will continue our effort to keep on expanding the catalog of cell types, adding further neuron attributes to the database and automatic techniques for classification of finding cell types based on structural, functional, transcriptional and physiological data. On the other hand, FishExplorer was designed to be extended directly by the community through authorized registration at the platform. Whether the community will actually engage into the platform needs to be seen. We will implement a programmatic interface via APIs to the databases stored in FishExplorer to enable the community to more easily access and analyze the data.

Finally, the field of neuroscience is witnessing a growing emphasis on comparative studies. Parallel and collaborative efforts form the backbone of this progress, and we recognize the importance of fostering a shared ecosystem for the teleost research community. To promote interoperability and cross-species analysis, we have recently integrated datasets from medaka (Vohra *et al.,* 2024) and are actively working to include additional species. For example, we plan to incorporate cellular-resolution data from electric fish (Sawtell et al. co-submitted with this manuscript), expanding the taxonomic breadth of the platform. In addition, to support studies of brain development, we will include datasets from different developmental stages. We envision FishExplorer as a continuously evolving, community-driven platform that will serve as a key resource for comparative neuroscience in teleosts.

FishExplorer represents a major step forward in zebrafish neuroscience by enabling users to bridge data types and scales – from brain-wide imaging to synapse-level anatomy – within a single, extensible platform. Researchers can trace neurons from EM datasets, localize them in light-level atlases, assess their activity profiles from functional imaging, and explore structure–function relationships across different modalities and even different datasets. Through its multimodal integration, FAIR-compliant design, and community-driven framework, FishExplorer facilitates scalable, reproducible circuit analysis and opens new avenues for mechanistic insight into brain function and behavior.

### Materials and Methods FishExplorer platform

FishExplorer and its integrated analytic tools are modular and use a microservices architecture to decouple data storage, visualization, and analysis modules. Each service is deployed separately on a custom cloud using Kubernetes. The scalable architecture allows us to add new services, replicate services, and independently update specific modules (e.g., updating the database, visualization and analysis functions for a new data type) without disrupting the entire system. All the services can be easily deployed to any other commercial cloud service provider, such as Google Cloud or Amazon Web Services. The frontend is component-driven, allowing for flexible mixing and matching of various components. The platform is cost effective and developed using standard open source components such as Babylon.js, D3.js, Vega and Vega-Lite (Satyanarayan *et al.,* 2017). The database component consists of a NoSQL database, namely MongoDB, which provides flexible schema and accommodates novel data types, while image stacks, models, meshes, and neuron morphologies are stored on a custom S3-like server (MinIO file server). The backend APIs and job services for access and automatic integration of user data in the platform are implemented in Python.

### Brain regions

MECE (Mutually Exclusive and Comprehensively Exhaustive) regions were generated by starting from a whole-brain mask encompassing the entire area covered by the original brain regions defined in Randlett et al., 2015. From this, a hierarchy starting with the major brain divisions (Level 1 = Forebrain, Midbrain, Hindbrain, Spinal Cord, Ganglia, Eyes) was generated. Overlaps between the original masks were removed and the regions were then linearly expanded to fill any empty space in the whole-brain masks. Borders were manually edited and smoothed using convolution. The same operations were applied to the subdivisions of each Level-1-region to generate Level 2 and beyond. For example, the Hindbrain was further subdivided based on the original masks for Rhombomeres 1 to 7, and subsequently Rhombomere 1 was subdivided into Cerebellum, Interpeduncular Nucleus, Locus Coeruleus, Oculomotor Nucleus nIV, Raphe - Superior, RoL-R1, and Ventrolateral Stripe of Serotonergic Neurons.

MISC (miscellaneous) regions were segmented using several marker-gene expression patterns, such as Serotonin (5-HT), Dopamine (TH) and Noradrenaline (NA). The organization and extensibility of these masks were detailed in a previous paper (Vohra *et al.,* 2024), which specifically addressed the platform’s functionality for cross-species comparisons.

### LM-EM bridges

The generation of LM-EM bridges is a complex problem. Petkova *et al., 2025* and Boulanger *et al.,* 2025 independently created LM-EM bridges for WBEM and FCLEM datasets, respectively. Here, we present the generic approach to create LM-EM bridges. As it is difficult to create such bridges using a very high-resolution EM stack due to the high cost of computation, a down-sized, low-resolution stack of the EM volume is usually used instead (the spatial resolution of EM volumes are described in Fig.2A and the resolution of LM is 0.798 × 0.798 × 2.00 um). Additionally, while preparing samples for EM imaging, researchers also collect other information, e.g., neurotransmitters, activity data, etc. using light-sheet imaging in the same animal. This kind of additional information can also be used in the registration process. There are two common strategies for creating LM-EM bridges:

1) Landmark-based: the registration is initiated by manually placing landmarks in both the LM reference brain and the low-resolution EM stack, iteratively adding or removing landmarks based on visual inspection until a satisfactory registration accuracy is achieved. In order to achieve reliable alignment, ∼10,000 landmarks are generally required for the whole brain (Petkova *et al.,* 2025).

2) Combination of automatic and landmark registration: a first bridge is generated between the LM reference brain and an additionally collected neurotransmitter or other confocal stack using a combination of linear, rigid and non-linear transformations. Second, another bridge is created by manually placing landmarks in the additionally collected confocal stack and the low-resolution EM stack, iteratively adding or removing landmarks based on visual inspection until satisfactory registration accuracy is achieved. The advantage of using extra information is that one can more easily see the one-one correspondence between neurons of the low-resolution EM stack and the additionally collected confocal stack, therefore placement of landmarks in this scenario is also easier. Finally, both transformations are combined to generate the final forward and inverse transformations, which can then be used to integrate EM datasets into FishExplorer

### LM-LM and LM-Xray bridges

A wide range of image- and point-based registrations was performed to establish bridges between datasets. Confocal registrations were carried out using the Advanced Normalisation Tools (ANTs) (Avant *et al.* 2007). We used the *T_AVG_antiERK12tERK* marker line to map mapZebrain (2019) to the FishExplorer reference brain. Both *T_AVG_antiERK12tERK* and FishExplorer reference stacks were preprocessed using histogram matching. Registration was then performed with rigid [-m MI, -c 1000×1000×1000×0, 1e-8,10, -f 12×8×4×2, -s 4×3×2×1], affine [-m MI, -c 1000×1000×1000×0, 1e-8,10, -f 12×8×4×2, -s 4×3×2×1] and SyN (diffeomorphic symmetric) [-m CC, -c 700×700×700×0, 1e-7,10, -f 12×8×4×2×1, -s 4×3×2×1×0] transformations, with the specified parameters. The registrations were carried out on a high-performance cluster with 300 GB memory and 60 CPU cores. Using the generated transformations, we mapped mapZebrain (2019) brain regions and neuron SWC files to the platform. Organ-level data (X-ray) and motor vagus (confocal stack) nerves were registered using BigWarp (Bogovic *et al.,* 2016). For this, we initiated registration by manually placing a few landmarks in both target and reference dataset, iteratively adding or removing points based on visual inspection until satisfactory registration accuracy was achieved. A list of tutorials including test datasets and step-by-step instructions for point- and image-based registrations with a Python script is available on the FishExplorer website https://zebrafishatlas.zib.de/tutorial/.

### Computation of custom NBLAST scoring matrix

The NBLAST algorithm (Costa *et al.,* 2016) creates pairwise similarity scores between a query and a target neuron based on morphology and position. Each neuron is divided into small segments, each is assigned a position and a vector describing its direction. For each segment in the query neuron, the closest segment in the target neuron is determined and compared with. The score should be high if two neurons belong to the same cell type and have similar structure, and low otherwise. This is achieved by empirically deriving the probability of a segment match or non-match from groups of neurons known to be matching or non-matching, and then creating a raw scoring matrix with the log probability ratio, depending on Euclidean distance between segment positions and the absolute dot product between the segment vectors.

We used the implementation in Python’s navis (Schlegel *et al.,* 2025) to create a new scoring matrix based on zebrafish neurons. To determine matching neurons for the zebrafish NBLAST scoring matrix, we used the fruit fly NBLAST scoring matrix to identify morphologically similar clusters within broader cell types. Using the fruit fly scoring matrix for very similar neurons in the zebrafish is expected to yield meaningful clusters (Costa et al., 2016). Groups of matching and non-matching neurons were defined based on cell types for mapZebrain from NeuroMorpho.Org (RRID:SCR_002145; Kunst *et al.,* 2019; Tecuatl *et al.,* 2024) and from FCLEM (Boulanger-Weill *et al.,* 2025). We used the four functional cell types – iMi, CMI, MON, and SMI – from Boulanger-Weill *et al.,* 2025, and accessed cell types from NeuroMorpho.Org via the corresponding API, and visually inspected each cell type. We chose four groups: Cerebellum eurydendroid (CE), retinal ganglion cells (RGC), subpallium principal cells (SPC), and medulla principal cells (MPC) as classes. Overall, this resulted in eight classes with potentially matching neurons, which were used to create eight folds.

#### Matching Neurons

Within each class, we wanted to find matching neurons that best predict morphological similarity. Since the FCLEM cell types are functionally defined and the cell types from NeuroMorpho.Org represent broader categories, we used the default NBLAST scoring matrix (based on the fruit fly neurons) to determine clusters of neurons with highest morphological similarity. We found that the functional FCLEM cell types are morphologically more distinct than subclusters within the mapZebrain cell types. Therefore, we matched each neuron from the FCLEM functional cell types only with its augmented version (as suggested in *navis* tutorials; Schlegel *et al.,* 2025) by drawing modifying parameters from a Gaussian distribution with mean 0 and standard deviation of 0.3 for translation, 0.005 for jittering individual coordinates, and 0.005 for scaling. This resulted in 78 groups of matching neurons, each containing two neurons. For the four mapZebrain cell types, we used hierarchical clustering with a dendrogram cutoff of 3.2 to identify subclusters within each type. The minimum mean NBLAST score per cluster for inclusion was -0.2 (note that these scores were generated with the fruit fly scoring matrix). As with the FCLEM neurons, we added the augmented version of each neuron to its matching group. This resulted in ten groups of matching neurons, with a mean (*M*) of 47.2, standard deviation (SD) of 26.28 neurons per group.

#### Non-matching Neurons

Non-matching neurons were also divided into eight folds. First, all neurons out of the eight classes that were not assigned as matching neurons were included as non-matching neurons for each fold. Then, all other neurons from the mapZebrain and FCLEM datasets were distributed equally among the eight folds (with any leftover neuron assigned to the last group).

#### Cross-validation

We then performed cross-validation over the eight folds, where the NBLAST scoring matrix for fold *i* was trained using all matching and non-matching neurons of all classes except class *i*. During testing, we evaluated performance across all eight folds (**Fig. 3B** for an example). We observed that, across folds, NBLAST raw scoring matrices were qualitatively similar and varied only slightly, for example in the smoothness of the value distribution. Consequently, the average scores across all eight folds for both matching and non-matching neurons are qualitatively similar. We selected the raw scoring matrix created in fold 6 for further evaluation, as it was the smoothest based on visual inspection.

### Hierarchical Clustering with NBLAST

For hierarchical clustering based on NBLAST scores, we followed the approach in the navis tutorials (Schlegel *et al.,* 2025), first running NBLAST between all *N* neurons to generate a matrix *S*. We then created a symmetric matrix *A=-((S+S.T)/2 -1)* to construct a dendrogram using Ward’s method and cluster based on height in the dendrogram (Python’s *scipy*; Müllner, 2011; Virtanen *et al.,* 2020).

### Animal ethics

All experiments followed institution IACUC protocols as determined by the Harvard University Faculty of Arts and Sciences standing committee on the use of animals in research and teaching. The animal experimentation protocols, 25-03 and 22-04 were submitted and approved by this institution’s animal care and use committee (IACUC). This institution has Animal Welfare Assurances on file with the Office for Laboratory Animal Welfare (OLAW), D16-00358 (A3593-01).

### Data access

The image, neurons and metadata used in the study are browsable in the FishExplorer platform (https://fishexplorer.zib.de/sandbox/). All the datasets will be available at our download page (https://fishexplorer.zib.de/sandbox/downloads).

### Github link

We believe in open source access therefore the source code of FishExplorer and CircuitExplorer will be available at https://github.com/zibneuro/fishexplorer

## Authors contributions

Conceptualization: SV, HCH, DB, FE, Analysis: SV, ME, JBW, Software: SV, ME, Visualization: SV, Writing original draft: SV, HCH, ME, YI, DB, Writing review & editing: SV, DB, HCH, YI, JBW, FE, Methodology: SV, ME, JBW, MP, YI, GFPS, KJH, FK, AB, OR, HCH, DB Resources: DB, FE, Experimental support: DB, FE, JWL, Funding acquisition: DB, FE, JWL, Supervision: DB, HCH, YI, FE

## Acknowledgements

We are grateful to the Engert lab at Harvard University for experimental and conceptual support. We thank the Herwig Baier lab at Max Planck Institute for sharing brain regions and neuron morphologies. This work was supported by the National Institute of Health (U19NS104653) and in part by the Deutsche Forschungsgemeinschaft (DFG). Florian Engert received funding from the National Institutes of Health (U19NS104653 and 1R01NS124017-01), DoD (W911NF2420112) and the Simons Foundation (SCGB 542973 and NC-GB-CULM-00003241-02). Armin Bahl was supported through the Emmy Noether Program (BA 5923/1-1), the Zukunftskolleg Konstanz, as well as the German Excellence Strategy (EXC 2117-422037984).

## Extended Data

**Extended Data Fig. 1.**
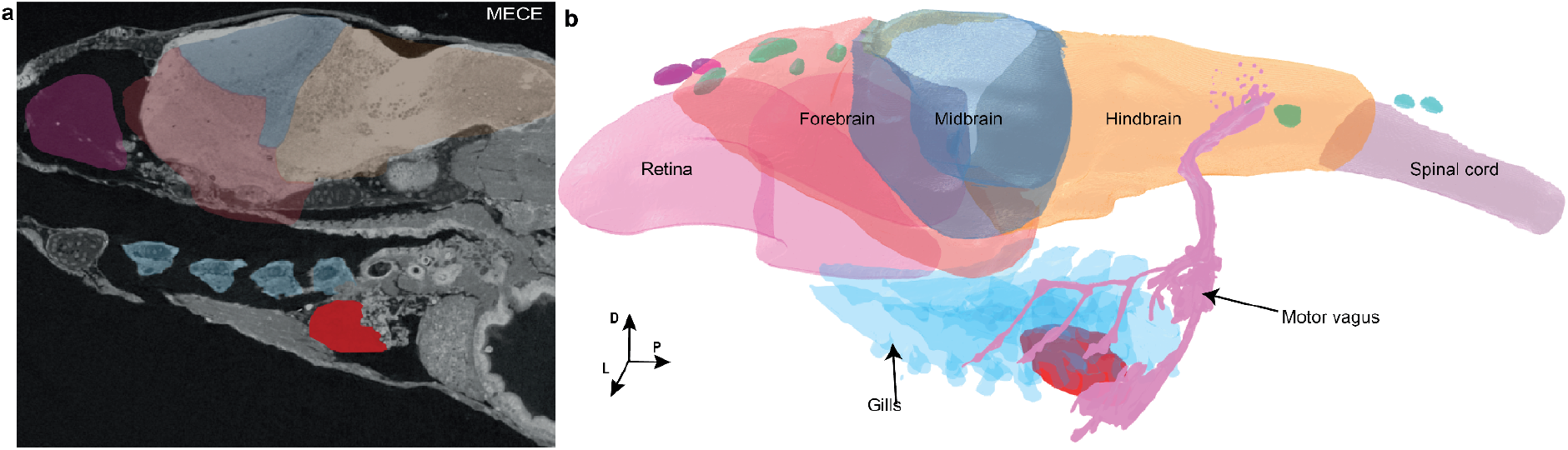
Body and the vagus nerve. **a,** A sagittal section shows an overlay of the vagus nerve and its motor elements including heart, gills and central motor nuclei. **b**, The heart, gills and other body regions were segmented using micro-CT x-ray volume and mapped onto the LM reference brain (MECE masks are shown).

**Extended Data Fig. 3.**
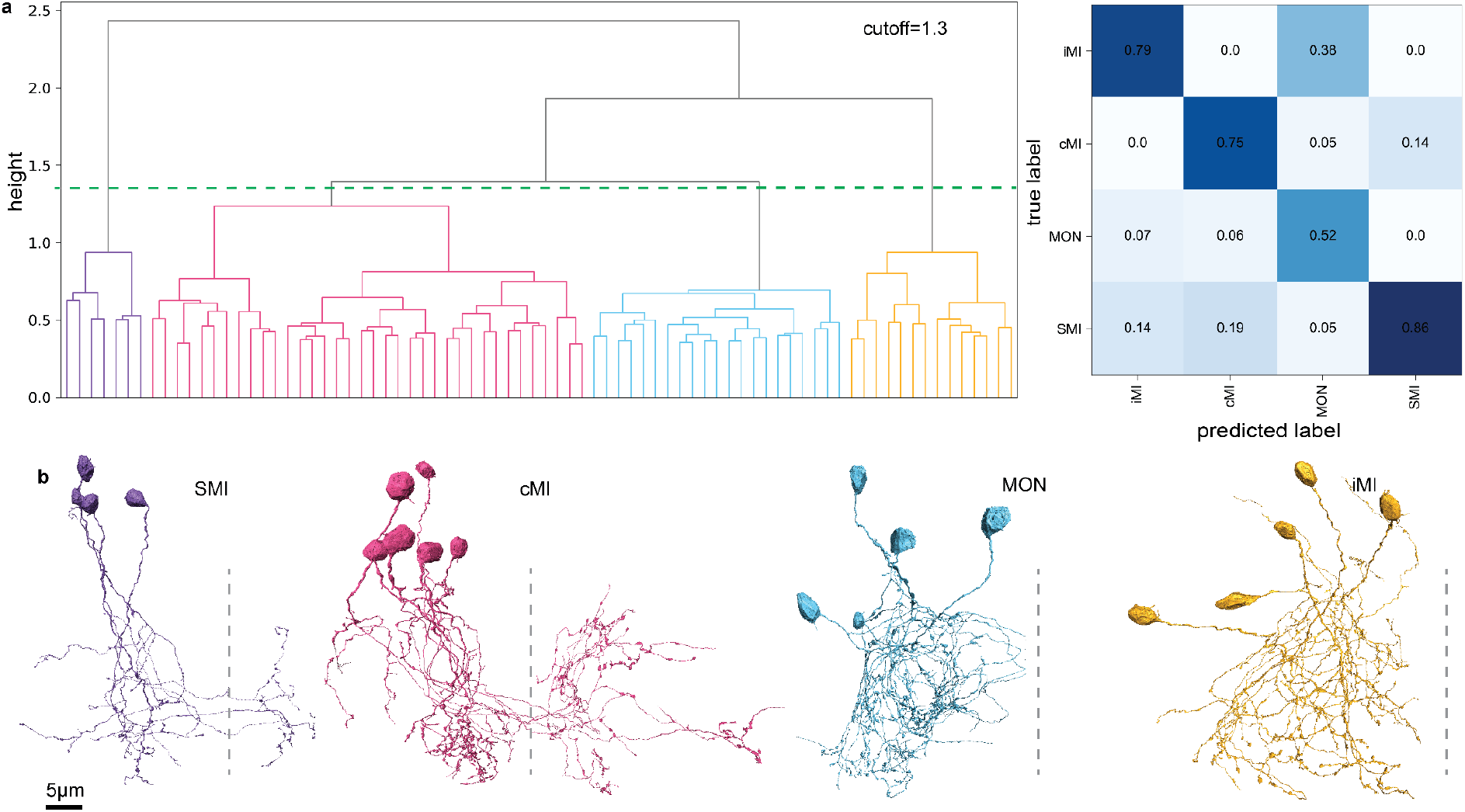
Hierarchical clustering and structural diversity within and between neuron clusters in FCLEM. **a**, Hierarchical clustering of functionally-labeled FCLEM neurons using a custom NBLAST matrix: The dendrogram on the left was cut at the height indicated by the dashed green line (cutoff = 1.3) to define distinct clusters, shown in different colors. The matrix on the right shows average NBLAST similarity scores within and between functional cell types. **b**, Visualization of predicted functional clusters: 3D rendering of neurons predicted to belong to iMI, cMI, MON and SMI clusters. ipsilateral motion integrator (iMi), contralateral projecting motion integrator (CMI), motion onset (MON), slow motion integrator (SMI). The dotted lines indicate the midline of the brain.

**Extended Data Table 1.** Meta-tags.

| Attribute label | Attribute description | Example |
| --- | --- | --- |
| Tracer's name | The name of the tracers those traced or update the cell | Petkova, Boulanger-Weill, etc |
| original data type | Name of the modality of the data | confocal, 2-photon, EM |
| Tracing date | Actual tracing date, followed by updating dates | 2024-04-28 |
| Neuroglancer cell ID | ID of the cell in the neuroglancer | 576460752330776649 |
| Neuroglancer soma ID | ID of the cell soma in the neuroglancer or cave | 576460752669123043 |
| Neuroglancer axon ID | ID of the cell axon in the neuroglancer | 576460752678666346 |
| Neuroglancer dendrite ID | ID of the cell dendrite in the neuroglancer or cave | 576460752659310512 |
| Publication | Name of the publications that the neuron is analyzed | Boulanger-Weill et al., |
| Neuron link | Neuroglancer or Cave link | https://spelunker.cave-explorer.org/#!middleauth+https://global.daf-apis.com/nglstate/api/v1/5624101328650240 |
| Lab | Name of the lab where the data was collected. | Engert |
| Molecular markers (if available ) | Name of a marker gene expressed in the neuron | Gad1b, vglut2a, etc |
| Functional labels (if available) | Name of functional cell type if the neuron is characterized | E.g.: integrator, motor command, etc |
| Other | For custom data | E.g.: percentage of reconstruction completed |

